# Remodelling of the glycome of B-cell precursor acute lymphoblastic leukemia cells developing drug-tolerance

**DOI:** 10.1101/2024.08.22.609211

**Authors:** Tiago Oliveira, Mingfeng Zhang, Chun-Wei Chen, Nicolle H. Packer, Mark von Itzstein, Nora Heisterkamp, Daniel Kolarich

**Affiliations:** Institute for Glycomics, Griffith University, Gold Coast Campus, QLD; Department of Systems Biology, Beckman Research Institute City of Hope, Monrovia, CA, USA; Department of Chemistry and Biomolecular Sciences, Macquarie University, Sydney; ARC Centre of Excellence for Nanoscale BioPhotonics, Griffith University, QLD and Macquarie University, NSW, Australia

**Author notes:** Institute of Molecular Biotechnology, Vienna Biocenter, Vienna, Austria (Tiago Oliveira). Corresponding authors, equal contribution: Nora Heisterkamp: Daniel Kolarich. <u>Email addresses:</u> Tiago Oliveira Mingfeng Zhang Chun-Wei Chen Nicole H. Packer Mark von Itzstein.

**Keywords:** human B-cell precursor acute lymphoblastic leukemia, drug resistance, minimal residual disease, vincristine, stromal co-culture, glycopeptide, HLA-DRA, LAMP1, bisecting N-glycans, MGAT3

## Abstract

Reduced responsiveness of precursor B-acute lymphoblastic leukemia (BCP-ALL) to chemotherapy can be first detected in the form of minimal residual disease leukemia cells that persist after 28 days of initial treatment. The ability of these cells to resist chemotherapy is partly due to the microenvironment of the bone marrow, which promotes leukemia cell growth and provides protection, particularly under these conditions of stress. It is unknown if and how the glycocalyx of such cells is remodelled during the development of tolerance to drug treatment, even though glycosylation is the most abundant cell surface post-translational modification present on the plasma membrane. To investigate this, we performed *omics* analysis of BCP-ALL cells that survived a 30-day vincristine chemotherapy treatment while in co-culture with bone marrow stromal cells. Proteomics showed decreased levels of some metabolic enzymes. Overall glycocalyx changes included a shift from Core-2 to less complex Core-1 O-glycans, and reduced overall sialylation, with a shift from α2-6 to α2-3 linked Neu5Ac. Interestingly, there was a clear increase in bisecting complex N-glycans with a concomitant increased mRNA expression of *MGAT3*, the only enzyme known to form bisecting N-glycans. These small but reproducible quantitative differences suggest that individual glycoproteins become differentially glycosylated. Glycoproteomics confirmed glycosite-specific modulation of cell surface and lysosomal proteins in drug-tolerant BCP-ALL cells, including HLA-DRA, CD38, LAMP1 and PPT1. We conclude that drug-tolerant persister leukemia cells that grow under continuous chemotherapy stress have characteristic glycotraits that correlate with and perhaps contribute to their ability to survive and could be tested as neoantigens in drug-resistant leukemia.

## Introduction

Long-term interactions between precursor B-acute lymphoblastic leukemia (BCP-ALL) cells and the bone marrow microenvironment involve the glycocalyx, the complex layer of glycoconjugates covering every cell. This is an essential communication component between the protective extracellular matrix, stromal cells, and immune cells that could attack the leukemia cells. Moreover, the glycocalyx structure is cell-type specific and can experience dynamic changes as a consequence of malignant transformation (1–6) but also potentially as a consequence of chemotherapy (7). To date, however, the glycome of chemotherapy-treated leukemia cells has received little attention. One previous study investigated N-glycosylated peptides, but compared a chronic myeloid leukemia K562 cell line that was fully resistant after long-term adriamycin (doxorubicin) treatment to a non-resistant cell line, without any stromal involvement (8). Also, in a CEM T-ALL model, the N-glycome of cells resistant and sensitive to the tubulin-binding agent epothilone B analogue 13-desoxyepothilone B were compared. In that study, a shift from α2-6 to α2-3 sialylation was identified on membrane glycoproteins in resistant cells which correlated with decreased expression of the *ST6Gal1* gene (7).

Thus, while these results do support the concept that chemotherapy induces changes in the leukemia cell glycocalyx, such studies do not comprehensively model the long-term process of environmentally-mediated drug resistance (EMDR) and drug-tolerant persistence (DTP) of leukemia cells: cells were exposed for a short period of time to drugs or were not provided with stromal support and cells were compared in which genetic alterations probably caused the drug resistance. To better investigate EMDR, we here employed a long-term co-culture system between human BCP-ALL cells and mitotically inactivated supporting murine stromal cells. We previously used this system to focus on the contribution of specific carbohydrate-binding proteins Galectin-1 and Galectin-3 to EMDR, and of a specific post-glycosylation modification, 9-*O*-acetylion of *N*-acetylneuraminic acid (9–12). Here, we analyzed glycomes of BCP-ALL cells able survive and grow in the long-term presence of vincristine. The combined results support the concept that specific glycocalyx alterations occur in cells that have become vincristine-resistant while receiving protection from the microenvironment. This may lead to identification of novel glycocalyx targets in BCP-ALL that developed vincristine resistance.

## Results

### Proteome of vincristine-resistant cells shows limited changes compared to controls

Non-leukemic healthy human pre-B cells proliferate and differentiate in the bone marrow, and are regulated by interactions with bone marrow stromal cells. Hence, *ex vivo* co-culture with bone marrow stromal cells can, to some extent, model the microenvironment of normal and malignant B-cell precursor cells (**Fig. S1A**). Such systems are widely used to support survival and growth of human BCP-ALL cells (9–16) and can be used to monitor drug treatment outcomes. We harvested cells for *-omics* analyses (**Fig. S1B**) from cultures treated continuously with 2 and 4 nM vincristine at a point when sufficient cells had accumulated with good viability, on d18 and d30 of culture, respectively. Results of RNA-seq analysis of these drug-tolerant persister (DTP) cells have been described (17).

Proteomics after HILIC fractionation identified 1153 proteins, with a median of 871 proteins per sample (**Table S1**). Of the 26 differentially expressed proteins in the LFQ analyses, 5 were increased after 2 nM vcr treatment (FDR≤5% and log2FC≥ 1) (**Fig 1A; Table S1**). Prolonged treatment with 4 nM vincristine revealed 101 differentially expressed proteins, of which 27 were found at higher levels in cells growing in the presence of vincristine (**Fig. 1B; Table S1**). Proteins with differential expression (DE) included 23 enzymes with signal transduction or metabolic function, as well as 31 proteins involved in DNA repair, transcription, and translation. Two well-known cell surface adhesion/signalling glycoproteins with abundant O-linked glycosylation on pre-B cells, CD43/SPN and CD45/PTPRC, exhibited decreased expression. Seven proteins involved in cytoskeletal organization also showed differential expression. Of these, gelsolin (GSN) was the only protein with significantly increased expression at d18 of 2 nM vincristine treatment (log fold difference of 1.05 and 1.65, **Fig. 1A**). A concomitant increase in *GSN* mRNA was also measured at d30 (17). Concordant downregulation of protein and mRNA levels were further found for the histone H1FX; CYCS, the mitochondrial cytochrome C involved in electron transport and initiation of apoptosis; three metabolic enzymes, LDHA, PFKP and PSAT1; and other proteins involved in regulation of energy and metabolism including GBE1, TPD52, TRIM72, and WARS, although transcriptomics and proteomics indicated different log-fold level changes of downregulation (**Fig. 1C**).

**Figure 1.**
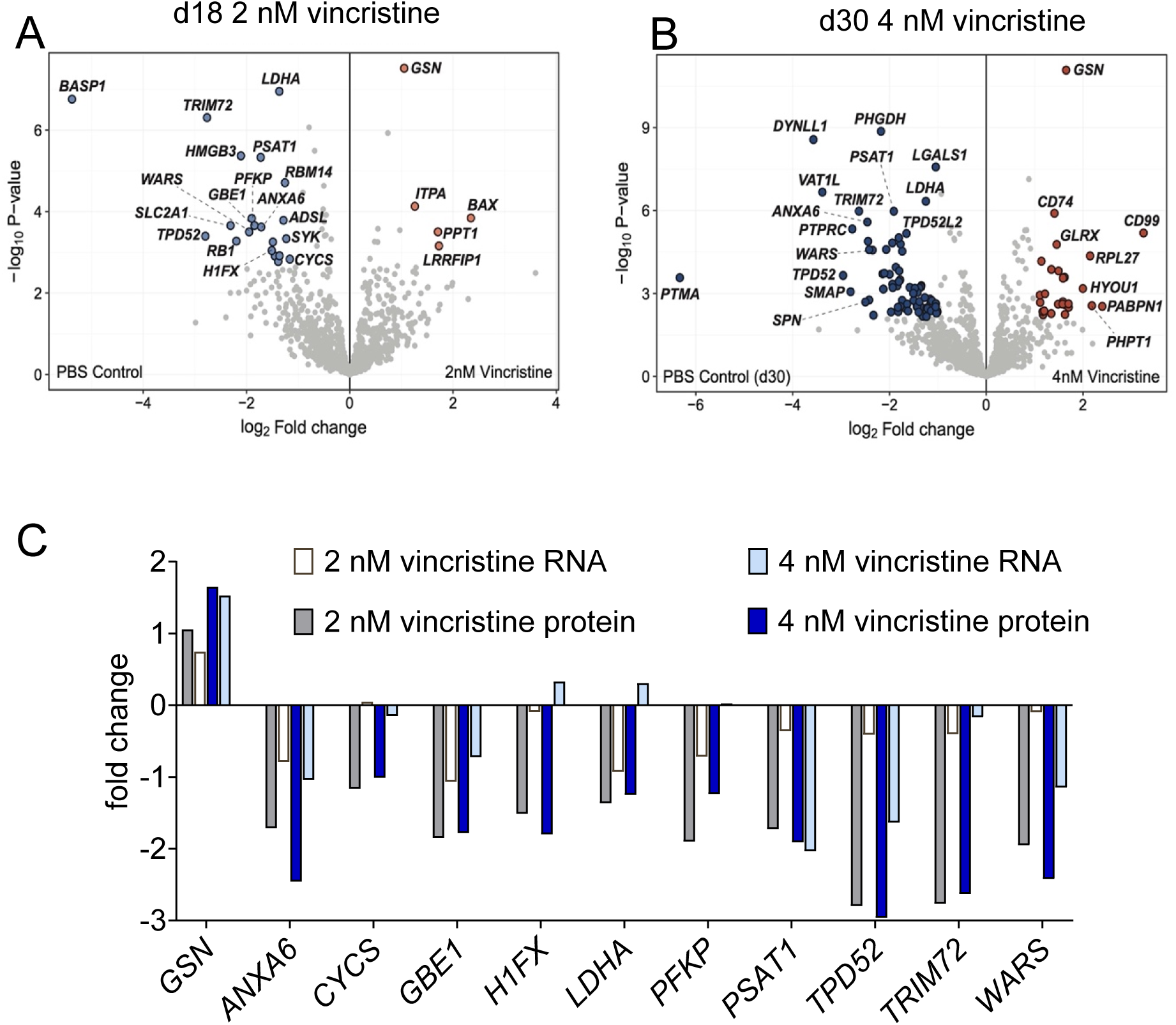
Analysis of the proteome of human ICN13 BCP-ALL cells after development of unresponsiveness to vincristine drug treatment in the presence of microenvironmental support. (**A**, **B**) Volcano plots of differential protein expression on d18 (26 DE proteins, **A**) and d30 (101 DE proteins, **B**). (**C**) Correlation of mRNA and protein expression of the 11 proteins that were found simultaneously differentially expressed when comparing both vincristine treatment conditions to the respective controls. ICN13 cells were co-cultured with mitotically inactivated OP9 stromal cells.

### Vincristine treatment leads to a modulation of the glycocalyx of BCP-ALL cells

These results combined with RNA-seq data (17) suggested a potential remodelling of the glycocalyx of leukemia cells growing in the presence of vincristine, as protein glycosylation enzymes are not only regulated at the transcriptional level, but also strongly influenced by cellular metabolic flux (18). We investigated this using well-established glycomics technologies to capture the N- and O-glycome as well as the proteoglycome contributed by glycosaminoglycans (GAGs). O-glycomics analyses identified 6 distinct structures of the Core 1 and Core 2 type (3 structures each, **Fig. 2A** and **Table S2**). The relative levels of Core 1 type O-glycans varied from 71 to 87% in all samples, while Core 2 O-glycans accounted for 12 to 29% of the overall glycome of these cells (**Fig. 2A**). No major change in the O-glycome was present in the d18, 2 nM treated cells, but the relative abundance of Core 1 type O-glycans was increased in the 4 nM vincristine-treated cells compared to controls, and accompanied by a reduction in Core 2 type O-glycans (**Fig. 2A**). No concomitant changes were evident at the level of mRNA transcripts of genes dedicated to the synthesis of O-glycans [*C1GALT1*; *GCNT1-3*], clearly demonstrating that changes at the transcript level do not automatically associate with alterations in the overall glycome of cells. Changes in the GAG composition were identified in cells after 30-days of exposure to vincristine compared to control cells, cultured in parallel without continuous vincristine exposure: non-sulphated heparan sulphate levels decreased, whereas those of hyaluronic acid, a major ligand of CD44, increased (**Fig. 2B** and **Table S3)**. Interestingly, previous results suggest that CD44 contributes to survival of BCP-ALL cells treated with vincristine (17).

**Figure 2.**
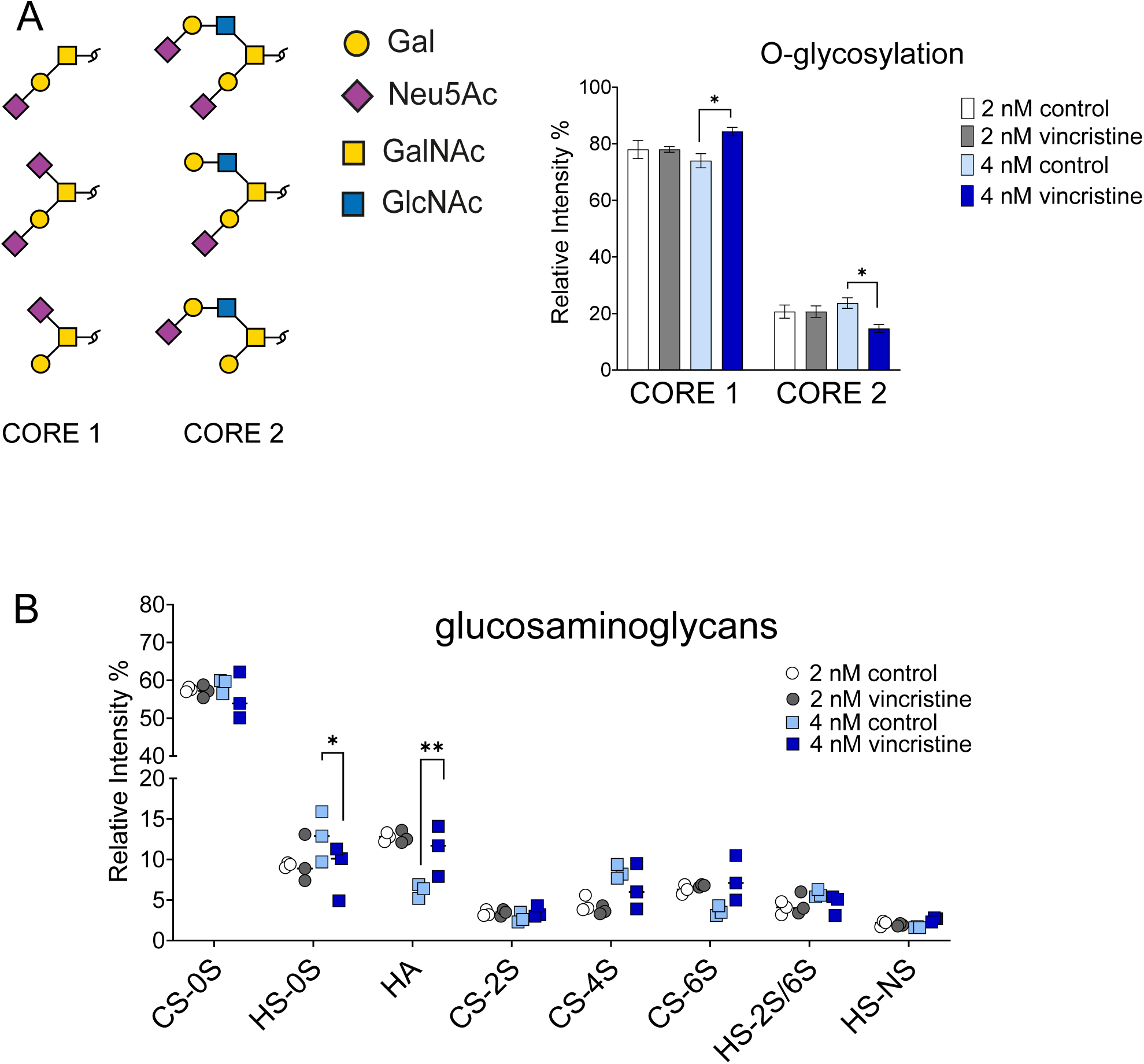
Complex O-linked glycan remodelling in BCP-ALL cell long-term survivors of vincristine treatment. (**A**) Relative representation of Core 1 and 2 type O-glycans in the indicated samples. The identified structures are represented on the left. (**B**) Relative abundance by fluorescence peak area quantitation of GAG disaccharide including CS, chondroitin sulphate; HS, heparan sulphate; and HA, hyaluronic acid. The various non- and mono-sulphated GAGs were reproducibly detected across all samples using 8 common HS and 8 common CS retention time standards and HS, CS, HA GAG digest control. HS-2S and HS-6S were not well resolved and quantified together. *p<0.05; **p<0.01

Overall levels of sialylation were reduced in the 4 nM vincristine cultured cells, in particular due to the decreased presence of α2-6 sialylated N-glycans (**Fig. 3A**), even though overall mRNA expression levels of *ST6GAL1* did not change (**Fig. 3B**). Mono-sialylated α2-3 and α2-3/α2-3 bi-sialylated structures N-glycans were increased, albeit not significantly (**Fig. 3A**). There was a significant increase in the mRNA expression levels of *ST3GAL5* [GM3 synthase](**Fig. 3C**). which has also been associated with drug resistance in other leukemia cell lines (19). Relative overall levels of fucosylation were variable between conditions. A 4.5% median overall decrease in fucosylated N-glycans was found for the d30 cells exposed to vincristine. This was due to a decrease in complex fucosylated structures, and paucimannose-type, core fucosylated N-glycans showed a slight increase (**Fig. 3D, E**).

**Figure 3.**
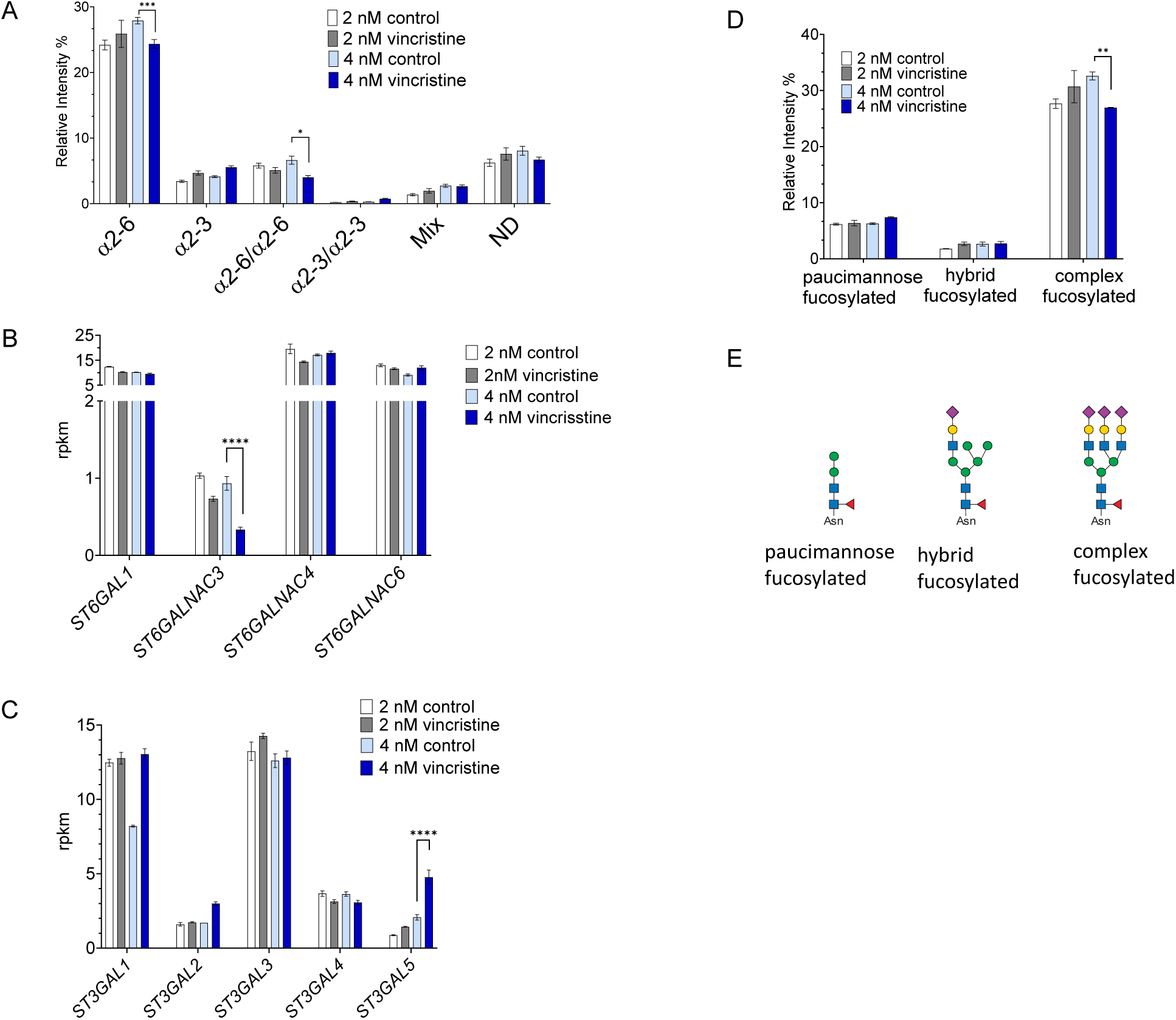
Long-term vincristine treatment correlates with overall decreased sialylation. (**A**) Relative abundance of N-glycan structures based on sialic acid linkage. (**B**, **C**) RNA expression of the indicated sialyltransferases. rpkm, reads per kilobase million. ***p<0.001 ****p<0.0001. (**D**) The relative abundance of fucosylated glycans in each group. (**E**) Graphical examples of fucosylated glycan structures. H, Hex; N, HexNAc; F, fucose; NAc, neuraminic acid.

We detected 75 distinct N-glycan structures distributed over four categories: 10 oligomannose, 7 paucimannose, 11 hybrid (of which 10 sialylated), and 47 complex type N-glycans, the majority of which were sialylated and/or fucosylated (**Fig. 4A, Table S4**). While the N-glycome remained largely stable, the 4 nM vincristine cultured cells presented the lowest level of complex sialylated glycans, and a median of 8.5% of paucimannosylated structures, which was approximately 1.5% more than the respective controls and all other sample groups (**Fig. 4A**). Interestingly, complex bisecting N-glycans accounted for 2% of the structures found on the vincristine cultured samples, whereas they were essentially absent in control cells (**Fig. 4A**). MGAT3 is the only glycosyltransferase that catalyses the formation of bisecting N-glycans within the medial Golgi apparatus (20–23). Notably, *MGAT3* transcript levels were upregulated in the cells chronically exposed to 4 nM vincristine (**Fig. 4B**). In fact, of the seven bisecting N-glycan structures, five were only present in the vincristine treated cells (**Fig. 4C**), confirming that *MGAT3* upregulation was associated with the increase of bisected N-glycans.

**Figure 4.**
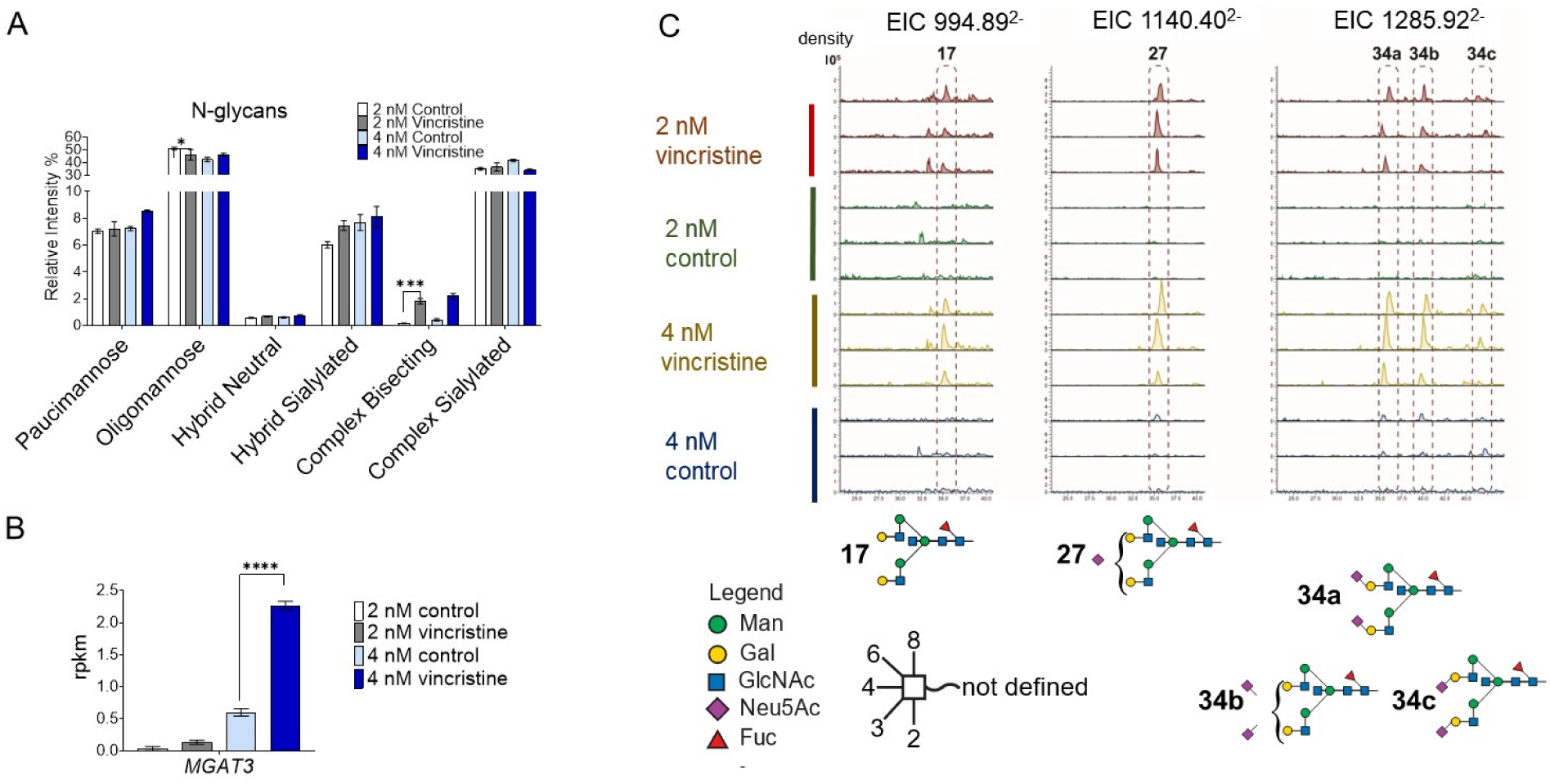
Increased levels of complex bisected N-glycans and increased *MGAT3* expression in BCP-ALL cells growing under chemotherapy treatment in the presence of stromal support. (**A**) Relative intensity of N-glycans in the major classes as indicated. (**B**) Expression of *MGAT3* transcripts induced by treatment with vincristine (**C**) EIC of m/z 994.89^2-^, 1140.40^2-^ and1285.92^2-^ demonstrate the differential expression of specific bisecting N-glycan structures indicated below the figure. Numbers refer to structures in **Table S4**. *p<0.05; ***p<0.001; ****p<0.0001

### Vincristine treatment induces distinct protein and site-specific changes in N-glycosylation

These results showed that subtle changes had occurred in the overall glycome of the DTP cells. Since cell surface glycoproteins are responsible for many of the interactions between leukemia cells, the extracellular matrix, and cellular components of the leukemia bone marrow microenvironment, we examined the glycoproteome of the leukemia cells. HILIC enrichment using microcrystalline cellulose, and subsequent analyses of intact glycopeptides allowed us to gather in-depth information about the most common glycosylation features on specific sites after vincristine treatment. We identified 114 distinct glycosites in 82 glycoproteins, including 1417 intact glycopeptides (**Table S5**), representing commonly expressed as well as B-cell precursor-characteristic proteins such as CD79a, CD79b and IGM. A limited number of O-glycan modified peptides were also detected including possible O-GlcNAc modification of HCFC1 and KDM3B (**Table 5**), known substrates of OGT (24, 25). The 82 individual glycoproteins originated from all subcellular compartments, with 28 mainly located at the plasma membrane or secreted, 11 lysosomal, 10 nuclear, 19 ER-resident or Golgi associated, and others with multiple locations. Some of these proteins were not identified in the initial proteomics analyses of the unfractionated samples, but only after HILIC fractionation. In addition, we verified their expression using the transcriptomics dataset of these cells (17).

Leukocyte Antigen class II histocompatibility antigen, DR alpha chain (HLA-DRA) plays a central role in the immune system by recognizing and presenting peptides derived from extracellular proteins (26). HLA-DRA showed a site-specific alteration of glycosylation in the 2 nM vincristine treated cells even though its overall levels were not significantly changed (**Fig. 5A, B**). 136 HLA-DRA associated glycopeptides were identified covering two sites of N-glycosylation, Asn103 and Asn143. We identified 130 glycopeptides that covered Asn103, which is highly conserved in evolution across mammals (27). The majority of glycopeptides carried largely complex type sialylated and/or fucosylated N-glycans in 9 distinct N-glycan compositions (**Fig. 5C**). The levels of some of these changed in response to chemotherapy and/or prolonged tissue culture (**Fig. 5C**). In the initial Byonic analyses, we were only able cover Asn143 (**Fig. S2**) in the vincristine treated cells, by identifying six glycopeptides carrying either Man_5_ or Man_6_ glycans. However, relative abundance analyses using Skyline did not detect any changes for these glycopeptides between conditions (**Fig. S2**).

**Figure 5.**
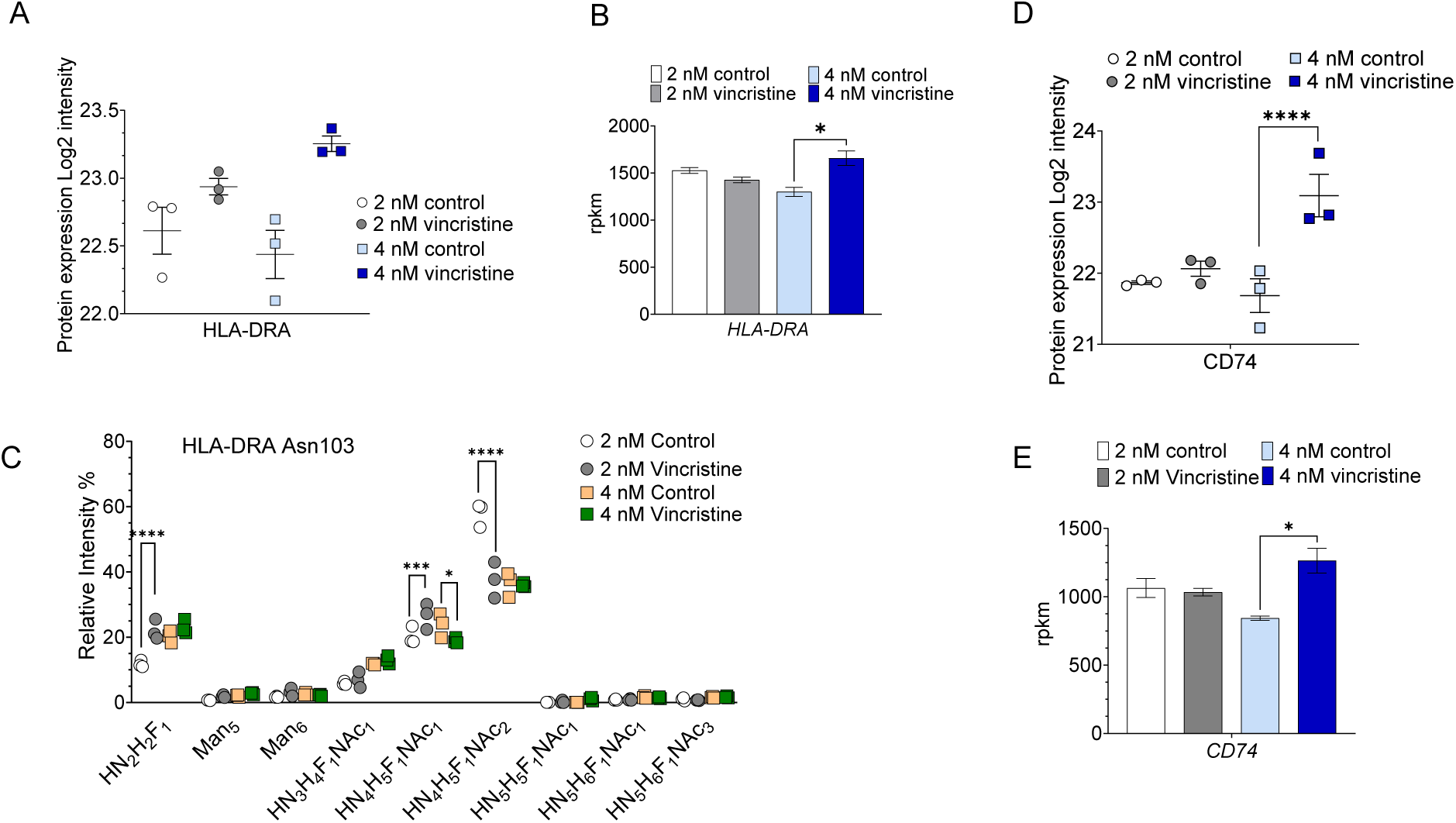
Altered expression and glycosylation of histocompatibility antigen components HLA-DRA and CD74 during chronic vincristine treatment. Expression levels of protein and mRNA of HLA-DRA (**A** and **B**) and CD74 (**D** and **E**). (**C**) Relative intensity of N-glycans attached to HLA-DRA Asn103. *p<0.05; ***p<0.001; ****p<0.0001

CD74 is also known as the HLA-DRA-associated invariant chain. We found a significant upregulation of CD74 at the protein level (**Fig. 5D**) and a moderate increase in corresponding mRNA in 4 nM vincristine treated cells levels (**Fig. 5E**). Glycoproteomic analyses yielded site-specific information for Asn130 and Asn136, two of the three previously described CD74 N-glycosylation sites. All 63 CD74 glycopeptides exclusively carried oligomannose type N-glycans, ranging from Man_5_ to Man_9_ (**Table S6**) with no significant changes in the DTP cells.

CD47 plays an important role in immune cell recognition, as adhesion receptor for THBS1 and as a “don’t-eat-me” signal for phagocytic cells (28). Four different CD47 N-glycan compositions were identified including three oligomannose, and one hybrid (**Fig. S3A**). Long-term vincristine treatment resulted in a small non-significant decrease in CD47 transcript levels. On the glycopeptide level, only oligomannose Man_7_ carrying N-glycans exhibited significantly reduced levels (**Fig. S3A**). We were not able to detect CD47 peptides in our proteomics analyses. Differential glycosylation of CD38, an extracellular cyclic ADP ribose hydrolase, will be described in more detail elsewhere.

Next to the cell surface, glycoproteins are also highly relevant constituents of the lumen-facing surface of lysosomal membranes (29, 30), where they play a major role in the protection of the lysosomal glycocalyx against degrading enzymes (31, 32). Several of the glycoproteins identified here have a lysosomal function including among others LAMP1, LAMP2, MPO, CTSA, CTSC, and PSAP. PPT1 (palmitoyl-protein thioesterase 1), **Fig. 6A, B**) breaks down lipid-modified proteins in the lysosome (33). Its protein expression was higher in 18-day 2 nM vincristine exposed cells compared to cells cultured for a similar period without drug treatment (**Fig. 6A**). Interestingly, the levels of paucimannose HexNAc_2_Hex_2_Fuc_1_ N-glycan attached to Asn197 almost doubled in vincristine treated cells (**Fig. 6B**).

**Figure 6.**
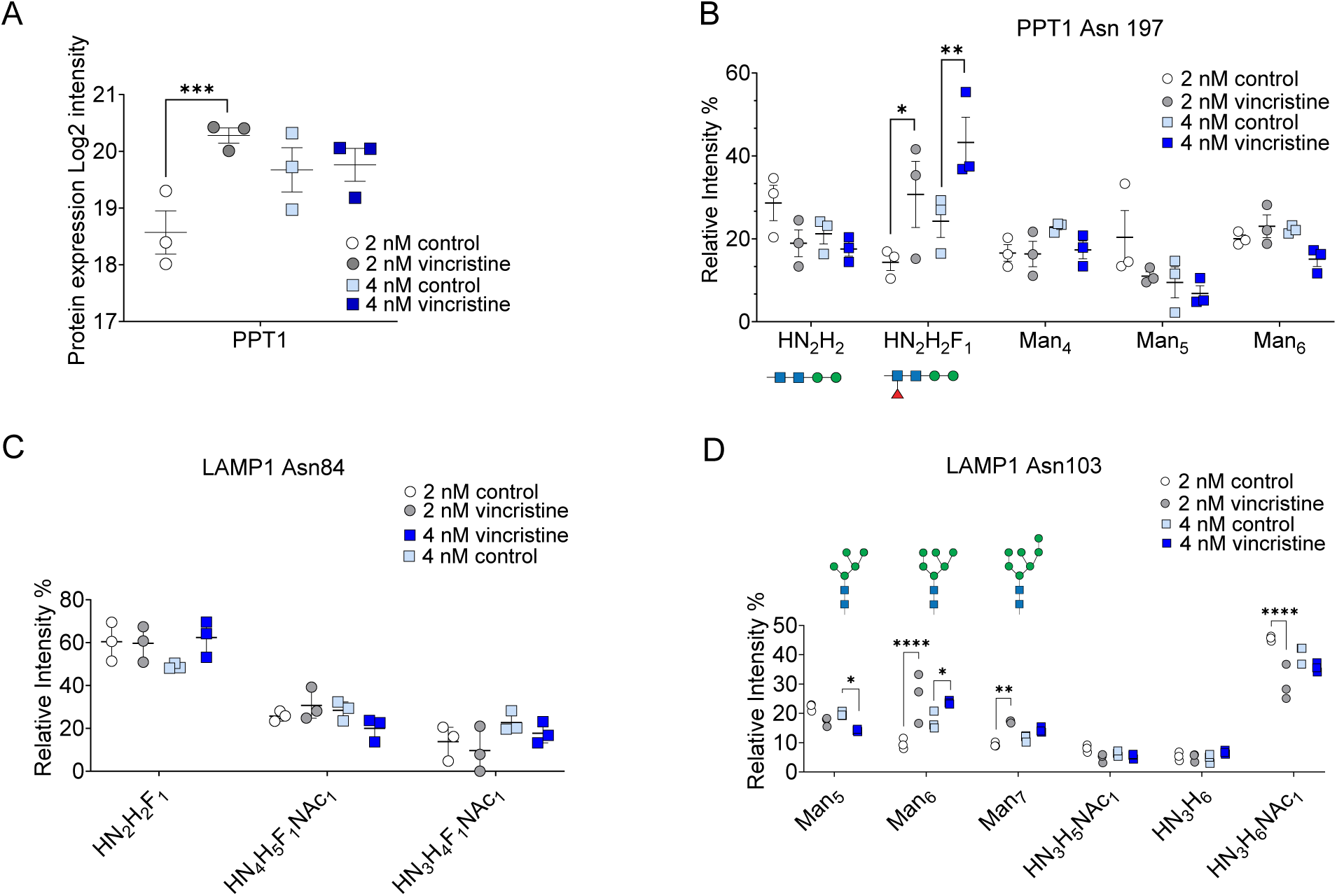
Evidence for remodelling of the lysosomal glycocalyx in BCP-ALL cells after 30 days of 4 nM vincristine chemotherapy. (**A**) Log2 protein expression of levels of PPT1 (**B**) Relative intensity of indicated glycan structures occupying Asn197 on PPT1. (**C, D**) Relative intensity of different glycan structures on LAMP1 Asn84 and Asn103.

Lysosome-associated membrane protein 1 (LAMP1) is important for the structural integrity of the lysosomal membrane. It contains a lysosomal lumen portion containing 18 potential sites of N-glycosylation (34). A single non-modified LAMP1 peptide (^299^FFLQGIQLNTILPDAR^314^) was repeatedly identified. A glycopeptide covering and carrying a Man_5_ oligomannose on Asn261 was identified in the PBS treated control samples. 128 glycopeptides covering N-glycosylation sites Asn84 (30 glycopeptides, **Fig. 6C**) and Asn103 (97 glycopeptides, **Fig. 6D**) were also detected. All samples contained the paucimannose N-glycan GlcNAc_2_Man_2_Fuc as predominant form of N-glycosylation on Asn84. Oligomannose type (Man_5_, Man_6_ or Man_7_) and hybrid, sialylated N-glycans (HexNAc_3_Hex_5_NeuAc and HexNAc_3_Hex_6_NeuAc) were the major forms of glycosylation occupying Asn103. Vincristine treatment resulted in increased levels of Man_6_ and Man_7_ N-glycans on LAMP1 Asn103 without a corresponding decrease in the expression of *MAN1A1* (**Fig. S5**) (35). The level of the sialylated hybrid structure HexNAc_3_Hex_6_NeuAc was significantly downregulated in the 2 nM vincristine-resistant cells when compared to controls (**Fig. 6D**).

### Cas9/CRISPR dropout screen confirms importance of glycosylation to the ability of BCP-ALL cells to survive vincristine treatment

We also tested 102 genes involved in glycan synthesis for their possible critical function in the ability of the KOPN8 BCP-ALL cell line to survive a 20-day, long-term vincristine treatment in co-culture with OP9 stroma cells (**Fig. 7A**). These genes were previously investigated for a possible non-redundant critical role in leukemia cell survival (36). Cells were harvested at a point when negative control sgRNA-containing cells had already largely been eliminated. Top ranking genes needed for overall survival had been previously identified as *OGT1*, *NGLY1*, *MGAT1* and *OGA* (36) and these remained important to survive long-term vincristine exposure. In addition, *C1GALT1C1*, *SLC33A1* and *B3GALNT2*, among others, ranked higher in contributing to survival during the 20-day vincristine treatment (**Fig. 7B, C**).

**Figure 7.**
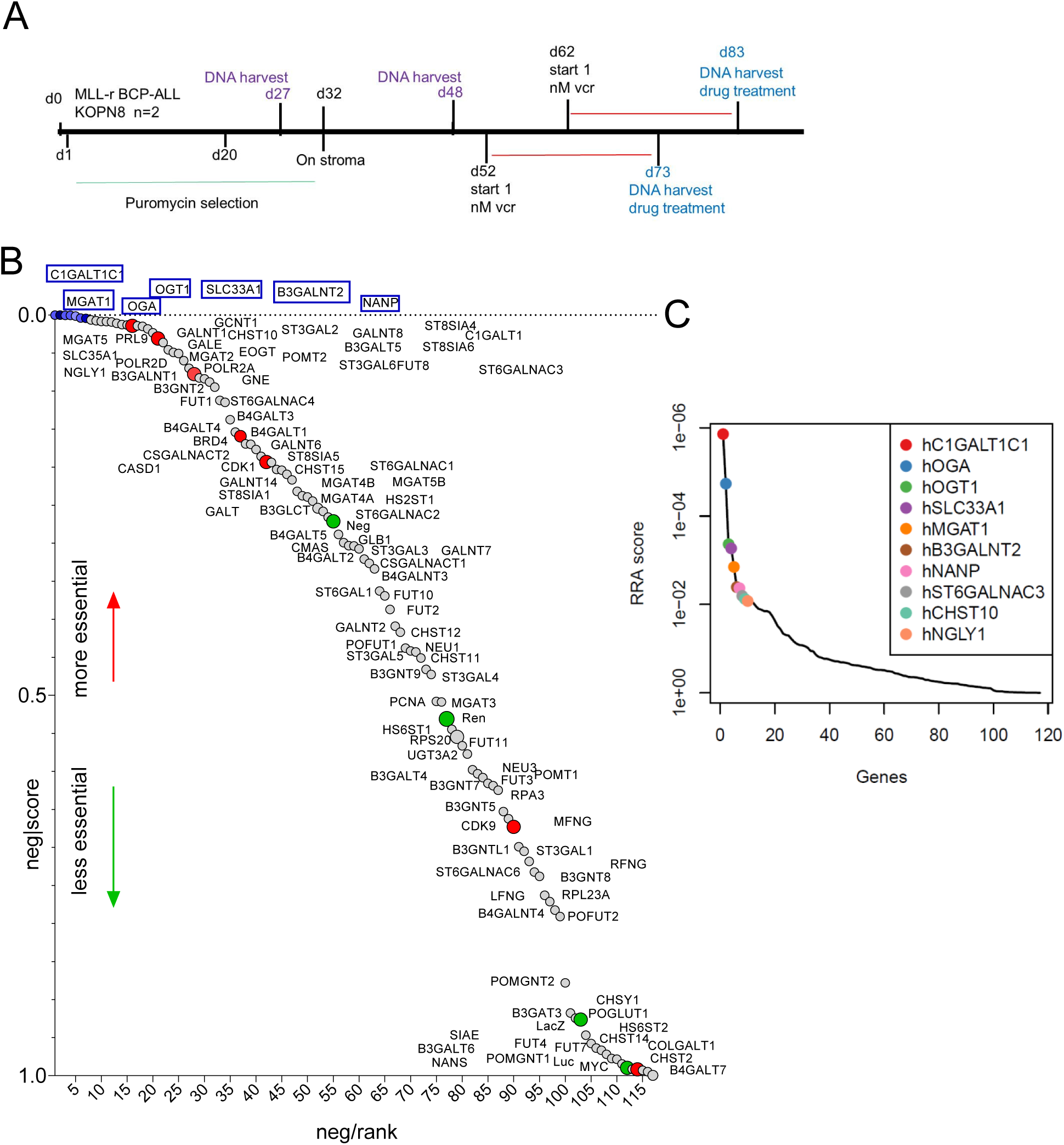
CRISPR screen to identify glyco-enzymes that support the emergence of DTP leukemia cells. (**A**) Schematic overview of treatment of KOPN8 BCP-ALL cells. Analysis of cells on d27 and d48 has been reported (36). (**B**) Negative-ranked MAGeCK score of glycan synthesis genes. Red dots, positive controls [essential genes]; green dots, neutral controls. (**C**) RRA scores ranking the top 10 genes contributing to persistence of KOPN8 cells chronically exposed to 1 nM vincristine.

## Discussion

### Changed proteome suggests modified adhesion

We previously noted changes in the RNA expression of genes regulating motility and adhesion in BCP-ALL cells that became resistant to vincristine (17). The proteomics data support this: we also measured decreased protein expression of both Annexin VI (ANXA6) and TPD52. ANXA6, a Ca^2+^-dependent, membrane binding protein, participates in membrane trafficking, membrane/cytoskeletal organization, and has been linked to cancer (37). It was reported as a candidate marker for monitoring minimal residual disease in BCP-ALL, with RNA levels upregulated more than 2-fold in patients (38). TPD52 is a candidate oncogene with increased expression in a variety of malignancies (39, 40). Overexpression of TPD52 was found to correlate with increases in ANXA6 levels and tumorigenesis (41). The simultaneous decrease in protein levels of both ANXA6 and TPD53 found in our studies in the cells cultured in the presence of vincristine further supports this connection.

CD99 is widely expressed on most haematopoietic cells, particularly on immature T-cells, and is involved in cell adhesion and migration, cell proliferation, differentiation, and apoptosis (42–44). High *CD99* mRNA levels are strongly associated with a higher frequency of relapse in pediatric BCP-ALL (45). This agrees with our results showing that drug-insensitive cells have significantly higher levels of CD99 mRNA and protein (**Fig. S4)**.

### Decreased contribution of Galectin-1

DTP BCP-ALL cells show hallmarks of increased quiescence (17). We here identified a shift from Core 2 to Core 1 type O-glycan representation in 4 nM vincristine-resistant cells when comparing to the respective control. This could be related to the decreased level of CD45 [PTPRC] and CD43 [SPN] in the d30 DTP BCP-ALL cells. CD45 and CD43 are major O-linked glycoproteins of which higher levels are associated with proliferation of immune cells and which are important ligands for Galectin-1 (46, 47)

Interestingly, proteomics revealed a significant decrease in Galectin-1 protein in the same cells (**Table S1**). This lectin is a modulator of immune response by binding to glyco-epitopes containing both LacNAc as well as α2-3 sialylated LacNAc present on both N- and O-glycans (48). Furthermore, Galectin-1 is a discriminating hallmark for the subcategory of BCP-ALL represented by the ICN13 cells used in the present study (11, 49). We found 135 MS/MS spectra corresponding to 3 distinct peptides covering 34.1% of the protein sequence of Galectin-1. Altogether, these results indicate that the glycome remodelling of 4 nM vincristine-resistant cells towards decreased Core 2 O-glycan expression and sialylation modulation occurs in parallel to a decrease in the expression of Galectin-1 protein, which suggests a downregulation of Galectin-1 involvement in drug-tolerant cells. This is consistent results from knockout studies of Galectin-1 in stromal cells showing it is dispensable for the development of drug insensitivity in BCP-ALL co-cultured cells (50).

### Bisected glycans show a significant increase

Vincristine-tolerant cells showed a remarkably increased representation of bisected N-glycans. MGAT3 is the only enzyme that catalyzes this branching (51), and we found a corresponding upregulation of *MGAT3* mRNA. MGAT3 activity is a key regulatory glycosylation step, because the bisecting GlcNAc it applies halts subsequent branching and terminal modifications (53, 54). In human adipose-derived stem cells and mature adipocytes, there was a significantly higher level of bisected N-glycans together with *MGAT3* mRNA (52).

The number of glycoproteins identified that are known to carry bisected N-glycans is increasing. Dan *et al* (55) identified 30 and 37 glycoproteins [with 17 in common] in two human cholangiocellular carcinoma cell lines that carry bisected glycans and these included LAMP1, LAMP2 and PSAP. In a normal human immortalized proximal tubular cell line HK-2, Dang *et al* (53) determined that 2% of N-glycans were bisecting N-glycans, and carrying proteins included among others HLA, PSAP, LAMP1/2, CTSC, TFRC and MME/CD10/neprilysin, which we also identified as glycoproteins in the ICN13 BCP-ALL cells (**Table S4**). They also reported that the expression levels of MGAT3 regulated glycan branching. Immunoglobulins such as IgG and IgM can also carry bisected N-glycans (56). Other known glycoproteins that are potential substrates of MGAT3 include the tetraspannin CD82 (57) and NOTCH1 (58).

However, because bisected N-glycans formed only a tiny subset of the total N-glycans, such structures were not recovered in our glycopeptide analysis and would require a focused effort to identify bisecting GlcNAc-carrying proteins, particularly if those proteins are not abundant. Interestingly, the expression of *MGAT3* is thought to be regulated by the transcription factors RUNX1 and RUNX3 (59, 60), and *RUNX3* mRNA levels were also increased in the vincristine-insensitive cells (17).

### Possible effects of glycan changes on lysosomal and/or immune functions

Our analysis of overall glycan composition showed that the emergence of DTP cells does not cause dramatic shifts in any of the major glycan categories. However, glycopeptide analysis showed that small quantitative differences measurable in the overall glycome do point to significant site-specific and protein-specific changes. This included proteins in the glycocalyx of both the cell surface and the lysosome, with possible interesting functional consequences.

PPT1, a lysosomal protein, is a molecular target of chloroquine and regulates autophagy (61, 62). Intact autophagy and lysosomal functions are clearly needed for the development of vincristine insensitivity of BCP-ALL cells (17). Seeing that glycosylation of N197 is essential for PPT1 to translocate to the lysosome (63), it is possible that specific glycan modifications can fine-tune its subcellular targeting and regulate autophagy. Interestingly, Hu *et al* (64) found significant increases in high mannose structures of lysosomal proteins in high-grade serous ovarian carcinoma cells compared to non-tumor tissues, supporting the concept that the lysosomal glycome can also be extensively remodelled during tumorgenesis.

Changes in CD74, HLA-DRA (**Fig. 5**) and CD47 (**Fig. S3**) glycosylation occurred when the BCP-ALL cells developed vincristine tolerance. Because these cell surface proteins mediate interactions with normal immune cells, their differential glycosylation could modulate the way the immune system recognizes and reacts to the leukemia cells. The effect of glycosylation on immune cell recognition was recently highlighted in Sharma *et al* (65). Using a mouse melanoma model, they showed that dendritic cells isolated from tumors have increased sialylation of PSAP. This leads to increased PSAP cell surface expression and reduced lysosomal location, where it would have been needed for processing of tumor antigens for cross-presentation to T-cells.

Thus, consistent with class II HLA expression on normal HSPC (66), ICN13 cells express HLA-DRA and could cross-present BCP-ALL tumor antigens to T-cells. However, in our analysis of BCP-ALL cells treated with vincristine, we did not recover sialylated PSAP (**Table S4**): although the number of unique glycopeptides identified was higher in the drug treated cells compared to controls [184 versus 32], N80, N101, N215 and N426 all carried HexNAc_2_Hex_2_Fuc_1_. This suggests that vincristine chemotherapy does not affect antigen processing for cross-presentation in these cells.

### Only two glycosyltransferases specifically promote BCP-ALL persistence during long-term vincristine treatment

Our Cas9/CRISPR dropout screen further showed that C1GALT1C1, SLC33A1 and B3GALNT2 expression is needed for the ability of the BCP-ALL cells to survive in the presence of vincristine. The need for SLC33A1 is not unexpected seeing that it transports acetyl-coA into the ER (67), regulates autophagy (68) and is essential as donor for acetylation of Neu5Ac (69). C1GALT1C1 is critical for O-glycan synthesis (70) and is needed for survival of T-cells outside the thymus in mice (71).

The reason ablation of B3GALNT2 gene function caused loss of fitness during vincristine treatment is less obvious. This enzyme synthesizes a very specific glycan structure, LacdiNAc (LDN; GalNAcβ1-3GlcNAc) and a limited number of 166 N-glycoproteins were identified that carry this structure (72) including NCSTN, which we previously identified as needed for survival of KOPN8 BCP-ALL cells (17). However, if LacdiNAc contributes to the structure and function of proteins that carry it, including NCSTN, it has not yet been reported.

### Glycopeptide analysis as a source of treatment targets

Intracellular leukemia-specific glycosylation, if induced by chemotherapy, could also be used as a treatment target. Malaker *et al* (73) detected O-GlcNAc peptides associated with MHC class I in leukemia samples which could function as neoantigens and possible treatment targets. They also identified O-GlcNac modified disaccharides attached to the same residues. Interestingly, in our glycopeptide analysis we found a number of potential O-GlcNAc or O-GalNAc modified intracellular proteins such as HCF1 and TLE4. Apart from HexNAc_1_, HCF1 also contained threonine-linked HexNAc_2_, whereas KDM3B, RAD23B, and PPP1R12A exclusively contained HexNAc_2_ (**Table 5**). Thus, future studies are warranted to determine if and how such glycosylation affects the function of some of these important glycoproteins in terms of BCP-ALL cell survival, and if differential glycosylation of intracellular or cell surface glycoproteins could be used as immunotherapy target.

## Acknowledgements

We thank Lu Yang for contributions to analysis of the CRISPR dropout screen results. Early studies were partly supported by a New Idea Award from the Leukemia Lymphoma Society and R01 CA090321 and CA172040 to NH, and a grant awarded to DK by Civic Solutions Inc. DK is the recipient of an Australian Research Council Future Fellowship (project number FT160100344) funded by the Australian Government. This work was made possible by a Griffith University Postgraduate Research Scholarship (GUPRS) awarded to TO.

## Materials and correspondence

Correspondence and requests for materials: Daniel Kolarich

## Supplemental data

This article contains supplemental data.

## Data availability

RNA-seq data have been previously reported and were deposited in GEO under number GSE176366.

## METHODS

### Reagents

Ultrapure water - Milli Q-8 system (Merck KGaA, Darmstadt, Germany) - was used in all experiments. All reagents were purchased from Sigma-Aldrich (St. Louis, MO, USA) at the highest quality, unless otherwise mentioned. PNGase F (Cat#P0705) was purchased from New England Biolabs (NEB). Sequencing-grade Trypsin from Roche (Cat#11047841001) was used for protein digestion. High-grade chloroform (Cat#1024442500) and methanol (Cat#1060072500) were purchased from Merck. A vincristine sulfate solution was obtained from Hospira Worldwide Inc. (Lake Forest, IL, USA).

### Cell culture and drugs

ICN13 PDX/patient-derived BCP-ALL cells and drug treatment are described in (17). Each biological replicate was divided into two fractions of 2x10^6^ cells, one for proteomic, glycoproteomic and glycomic analyses, and the second for RNA sequencing. Samples were stored at -80°C until analysis.

### Sample preparation for (glyco)protein extraction

For each biological replicate, 2x10^6^ BCP-ALL cells were lysed using a buffer composed of Dulbecco’s PBS (Sigma, Cat#D8537), 1% Triton X-100 (Sigma, Cat#T8787), Protease Inhibitors (Sigma, Cat#P8340) and Phosphatase inhibitors (Cell Signaling Technology, Cat#5870S). 200 µL of the cell lysis buffer were added to the cells, and samples were kept on ice for 20 min before being homogenised using a T-10 ULTRA-TURRAX® (IKA, Cat#0003737000) by performing 3 x 3 sec cycles (30 sec on ice between cycles). Samples were incubated on ice for another 10 minutes before a 30 sec water bath sonication. Cellular debris was pelleted by centrifugation for 15 min at 14,000xg, 4°C, and the resulting supernatants were collected to new tubes. All aliquots were stored at -80°C prior use. Protein concentration was estimated using a Pierce™ BCA Protein Assay Kit (Thermo Scientific, Cat#23225).

### N- and O-glycan release and PGC-LC-ESI-MS/MS glycomics

For each biological replicate, 100 µg of (glyco)proteins were reduced by adding 20 mM DTT, at 50°C for 30 minutes, and subsequently alkylated using 40 mM Iodoacetamide (at room temperature, in the dark). The reaction was stopped after 1 hour by adding another 20 mM of DTT and incubating for 5 minutes.

Proteins were precipitated as described previously using the chloroform:methanol:water separation (74, 75). The final protein pellet was air-dried for 15 minutes, before being resuspended using 10 µL of 8 M Urea and extensive vortexing. Glycoproteins were dot blotted (two dots each) onto a PVDF membrane (Immobilon P, 0.22 µm pore, Merck Millipore), and N- and O-glycans were sequentially released, as previously described (76). Briefly, N-glycan release was performed using PNGase F (50 U in MQ-H_2_O) at 37°C overnight, with the resulting N-glycans being collected to a new tube to be further processed and cleaned. O-glycans were subsequently chemically released from the same original dots by reductive β-elimination reaction. Before LC-MS analyses, reduced N- and O-glycans were desalted by cation exchange chromatography (AG50W-X8 cation exchange resin; BioRad, Cat#142-1431) (76, 77). To avoid the introduction of any potential contaminants in the LC-MS system, glycans were carbon cleaned using porous-graphitised carbon (PGC) material packed on top of C18 ZipTips (76). The N- and O-glycans were then stored at -20°C until being analysed.

The N- and O-glycome was determined using the PGC-nanoLC-ESI-MS/MS glycomics technology as described previously (76, 78). All details are described following the respective MIRAGE (Minimum Information Required for A Glycomics Experiment) guidelines in the Supplementary Material (79, 80).

### Glycan structure determination and relative quantitation

Glycan structures were determined as described previously (75, 77, 78). The registered *m/z* were manually analysed to determine each glycan structure, taking into account the observed PGC column relative retention time of each compound (76, 81–85). Glycan analyses was done with the help of GlycoMod (86), GlycoWorkBench software (v2.1) (87) and UniCarb-DB (https://unicarb-db.expasy.org/) (88). The most relevant signals present in the individual product ion spectra were used to assign a glycan structure, as described in “Comment-Diagnostic ions (*m/z*)” (**Tables S4** and **S5**).

Relative quantitation was performed for each determined compound present in any extracted ion chromatogram (EICs) of the corresponding monoisotopic precursors (83, 84). Spectra visualisation and area under the curve (AUC) acquisition were performed manually using Compass Data Analysis v4.2 SR1 (Bruker, Bremen, Germany), after smoothing the chromatograms (Gauss algorithm, 1s, 1 cycle). The AUC values of all the different detected/quantified glycans in the same sample were added (value corresponding to 100%) and the relative abundance determined for the corresponding AUC value. The calculated relative intensities were plotted using GraphPad Prism (v8.4.3), and *p*-values were calculated by performing an unpaired *t*-test. Symbols of calculated significance (*p*<0.05, **\***) are represented when groups are significantly different.

### Glycosaminoglycan disaccharide labelling

For GAG disaccharide profiling, 25 μg of (glyco)proteins of each biological replicate were dot blotted on to PVDF, following the same procedure as for N- and O-glycan analyses. An enzyme mixture (20 μL) containing 5 mU chondroitinase ABC from *Proteus vulgaris* (Sigma Aldrich), 2.5 mU heparinase I, II and III from *Flavobacterium heparinum* (Ibex Pharmaceuticals) in 100 mM ammonium acetate, pH 7 with 5 mM CaCl_2_ was added directly to each sample for incubation at 37°C for 16 hours. A glycosaminoglycan mixture containing heparan sulphate from bovine kidney, chondroitin sulphate from bovine trachea and hyaluronic acid from *Streptococcus* (all from Sigma Aldrich) was digested in parallel as an enzyme activity control, and as a retention time standard for hyaluronic acid. Released disaccharides were dried under vacuum, before being resuspended in 10 μL of water and labelled with 2-aminobenzamide (2-AB) according to manufacturer protocol (Ludger, Cat#LT-KAB-VP24-Guide-v2.0), using 2-picoline borane complex as a reducing agent. Excess 2-AB was removed according to Chu et. al. (89), using octanal as a water insoluble aldehyde reactant to conjugate excess 2-AB. In brief, the labelling reaction was topped up to 100 μL with deionized water and 400 μL of octanal was added and vigorously mixed. Phase separation was aided by centrifugation (60 s at 10000 rpm). The octanal layer was carefully removed, and 80 μL of the bottom aqueous layer was carefully removed, dried under vacuum and reconstituted in 32 μL of 75% (v/v) acetonitrile with 10 mM ammonium acetate pH 6.8; 4 μL (10% total sample equivalent) was injected onto the ZIC-HILIC column.

### ZIC-HILIC separation of 2-AB GAG disaccharide

ZIC-HILIC chromatography was carried out according to Takegawa et. al. (90), with some modifications. Chromatography was performed using Sequant ZIC-HILIC column (1 mm x 150 mm, 3.5 μm, 200 Å, Merck) on an Agilent 1260 Infinity HPLC at 50 μL/min flowrate in normal flow mode, with column oven set at 35°C, with fluorescence detection of the 2-AB label at 330 nm excitation/420 nm emission. The gradient used was as follows: Buffer A: 10 mM ammonium acetate pH 6.8, Buffer B: 90% acetonitrile with 10 mM ammonium acetate pH 6.8, 0 min – 100% B, 2 min – 90% B, 45 min – 30% B, 48 min – 10% B, 60 min – 0% B. To re-equilibrate the column, a short 8 min isocratic method of 100% B at 20 μl/min in microflow mode was used to trigger the fast-compositional change of the HPLC system. A mixture containing the 8 common heparin sulphate (Heparin disaccharide mix, Iduron) and the 8 common chondroitin sulphate disaccharides (Chondroitin/Dermatan sulphate disaccharide mix, Iduron) were used for the identification of disaccharides by retention time. Retention time of hyaluronic acid disaccharide was determined from the glycosaminoglycan mixture. Fluorescence peak area of each disaccharide was calculated using Agilent Openlab software.

### HILIC fractionation for glycopeptide enrichment and analyses

50 µg of protein was reduced, alkylated and precipitated following the aforementioned described chloroform-methanol protocol. The protein pellet was air-dried for 15 min before adding 100 µL of 25 mM ammonium bicarbonate prepared in MQ-H_2_O. 2 µg of High-grade Trypsin was added to each tube and samples were incubated for 18 h at 37°C. Peptides were then dried under vacuum and stored at -80°C until being used.

Hydrophilic interaction liquid chromatography (HILIC) was performed as described with a few modifications (91). Briefly, (glyco)peptides were resuspended in 20 µL of 1% TFA solution in MQ-H_2_O and extensively vortexed for 10 seconds. 80 µL of ACN/1% TFA was added very slowly to each tube to avoid precipitation of peptides. Pre-conditioned microcrystalline cellulose (50 µm, Sigma, Cat#435236) in 80%ACN/1% TFA was added to each tube. Tubes were incubated for 15 minutes in a rotor at room temperature before centrifuging the sample at 10,000 x *g* for 3 min to separate the cellulose from the solvent. The supernatant was carefully removed from the microcrystalline cellulose pellet and collected in a fresh Eppendorf tube. The pellet was washed three times with 100 µL of 80% ACN/1% TFA and the resulting supernatants combined with the first (flowthrough fraction). (Glyco)peptides were then eluted by adding 100 µL of 1% TFA to the pellet and incubating it for 5 min in the rotor at room temperature, followed by centrifugation and collection of supernatant. This step was repeated twice (elution fraction). The resulting fractions were cleaned by performing an offline C18 clean-up. The resulting C18 elution samples were dried under vacuum before being analysed by nanoLC-ESI-MS/MS.

The off-line HILIC fractionated peptides were identified using an Orbitrap Fusion™ Tribrid™ Mass Spectrometer coupled to an UltiMate™ 3000 UHPLC nanoLC (both Thermo Scientific™). Based on the NanoDrop quantitation, a volume corresponding to 600 ng of peptides were injected of each fraction. A MonoCap C18 Trap column (0.2x50 mm, 3 μm particle sizes, 100 Å pore sizes, GL Sciences, Cat#5020-10033) was used as a pre-column, whilst a MonoCap C18 Nano Flow (0.1x150 mm, 3 μm particle sizes, 100 Å pore sizes, GL Sciences, Cat#5020-10151) column was used for separation. The nanoLC resolving gradient was performed over 76 minutes for the separation of the peptides and subsequent column re-equilibration was performed over 44 minutes. Solvent A consisted of 0.1% TFA; solvent B was 90% ACN/0.1% TFA. The flow rate was held at 0.5 µL/min on the separation column. Sample loading onto the trap column was achieved within 6.5 min at 1% solvent B and a flowrate of 6 µL/min. The valve switch bringing the precolumn in line with the separation column was initiated at 6.5 min and a separation gradient was started as follows: increase of solvent B from 1-15% over 3 min, followed by an increase from 15-30% over 50 min, then 30-35% over 15 min, 35-45% over 5 min, and finally 45-90% over 3 min. The separation column was then re-equilibrated using following gradient of solvent B: 90% over 5 min, 90%-1% over 2 min, 1% over 30 min.

The Orbitrap Fusion mass spectrometer was operated in positive polarity mode with the Ion Transfer Tube temperature maintained at 300°C. The cycle time allowed for a full MS scan and subsequent MS/MS events was 3 sec. The full MS scan was performed at a 60K resolution in the Orbitrap with an automatic gain control (AGC) target value of 1x10⁶ (normalised AGC = 250%) for a scan range between *m/z* 600-1800. Charge states between 2-6 were allowed. Dynamic exclusion was enabled for 30 sec after 1 time, with +/-10 ppm. Precursors were prioritised based on their higher charge states. Before fragmentation, precursors were subjected to Quadrupole isolation using *m/z* 2 narrow isolation window, maximum injection time 250 ms. MS/MS scan first mass was defined as *m/z* 110. Higher-energy collisional dissociation (HCD) spectra were obtained in Profile mode at 30K resolution in the Orbitrap with an AGC target value of 5x10⁵ (normalised AGC=1000%), following assisted HCD (aHCD) fragmentation using 20,35,50% collision energies.

Glycopeptide identification was performed using Byonic (v2.16.11, Protein Metrics Inc.) on the *raw* data files corresponding to the HILIC elution fractions. The HCD-product ion spectra were searched against *in silico* tryptic digest of *Homo sapiens* proteins from the UniProt sequence database (reviewed sequences v10; May 2020), allowing up to two missed cleavages. The following modifications were used: cysteine carbamidomethylation was set as a fixed modification; methionine oxidation, glycan modifications, and acetylation of protein N-terminal were allowed as variable modifications. The glycan search settings were as follows: allowing Asparagine (Asn) modification for N-glycans, and Serine (Ser) and Threonine (Thr) modification for O-glycans. “N-glycan 309 mammalian no sodium” and “O-glycan 78 mammalian” databases were used as glycosylation parameters. Glycopeptides were initially filtered using a Byonic score >250. The results were then manually validated to avoid glycopeptides identified solely once, or without any corresponding MaxQuant Proteomics data.

We then integrated the area under the curve of these detected glycopeptides using the Skyline© software (v20.2.0.343). Skyline© required the detected retention time, charge state and *m/z* of these glycopeptides in order to perform the calculation of the AUC. The filtered output of Byonic was used to create a transition list imported into Skyline, and all the raw files of the “HILIC elution” fractions were used to perform the calculations, with the integration limits being manually validated. The relative abundance of the glycan on a particular position was estimated similarly to the procedure used for glycan analyses. Briefly, for each analysed site, the AUC of the detected modified glycopeptides was added, corresponding to 100%. For example, in the case of CD47 we found 4 possible glycan compositions attached to Asn73; therefore, for any given sample, the AUCs of these 4 compositions were summed corresponding to 100%. The relative abundance was finally calculated for each respective AUC.

### Proteomics data analyses

HILIC Elution and Flowthrough fraction files were analysed using the Andromeda search engine integrated into the MaxQuant suit (v6.10.43)(92). For each biological replicate, the fractions were combined according to their respective sample and injection. As two injections were made for each fraction, this resulted in two combinations of 2 fractions for each sample (injection 1 and injection 2). The HCD-MS/MS spectra were searched against *in silico* tryptic digest of *Homo sapiens* proteins from the UniProt sequence database (reviewed sequences v10; May 2020) containing 20,359 reviewed protein sequences (Swiss-Prot IDs). All MS/MS spectra were searched with the pre-set MaxQuant parameters, and the following modifications were used: cysteine carbamidomethylation was set as a fixed modification; methionine oxidation, acetylation of protein N-terminus, and asparagine deamination were allowed as variable modifications. False discovery rate (FDR) of the peptide spectral matches (PSMs), protein, and site were set to 1% based on Andromeda score. Match between runs (MBR) algorithm was activated to allow matching MS features between the different sample fractions and improve quantification (92).

LFQ-Analyst was used for label-free quantitation (LFQ) of the MaxQuant pre-processed proteomic datasets (93). Four main “Conditions” were defined as “PBS d18”, “PBS d30”, “2 nM Vcr” and “4 nM Vcr”, and each injection was used as an independent replicate. In the Advanced Options setting, the “Adjusted *p*-value cut-off” was defined to 0.01 (q-value, FDR<1%), whilst the “Log2 fold change cut-off” (log2FC) was defined to 2. All differentially expressed proteins were manually validated. GraphPad Prism (v8.4.3) was used for the RNA^-^protein integrative analyses, protein analyses as well as for the representation of the gene expression datasets.

### Experimental Design and Statistical Rationale

Each of the triplicate samples was grown in a separate well on OP9 stromal cells from the same batch of mitotic inactivation. To ensure consistency, vincristine and PBS control cultures were grown in parallel within the same experiment. Statistical rationale was as described previously, for each individual analyses (17, 36).

## Submission declaration

The work described has not been published previously, is not under consideration for publication; it is approved by all authors and tacitly or explicitly by the responsible authorities where the work was carried out.

## Author contributions

**Tiago Oliveira:** conceptualization, data curation, formal analysis, investigation; methodology, writing drafts

**Mingfeng Zhang:** cconceptualization, data curation, formal analysis, investigation; methodology

**Chun-Wei Chen:** conceptualization, methodology, reagents

**Nicole H. Packer:** conceptualization, review

**Mark von Itzstein:** funding acquisition, conceptualization, review

**Nora Heisterkamp:** conceptualization, data curation, formal analysis, funding acquisition, investigation; methodology, supervision, writing drafts, review and editing

**Daniel Kolarich:** conceptualization, data curation, formal analysis, funding acquisition, investigation; methodology, supervision, review

## Abbreviations

DTP: drug-tolerant persister
EMDR: environment-mediated drug resistance
GAG: glycosaminoglycan
logFc: log2-fold change
BCP-ALL: precursor B-cell acute lymphoblastic leukemia

## Supplementary Figures S1-S5

**Figure S1.**
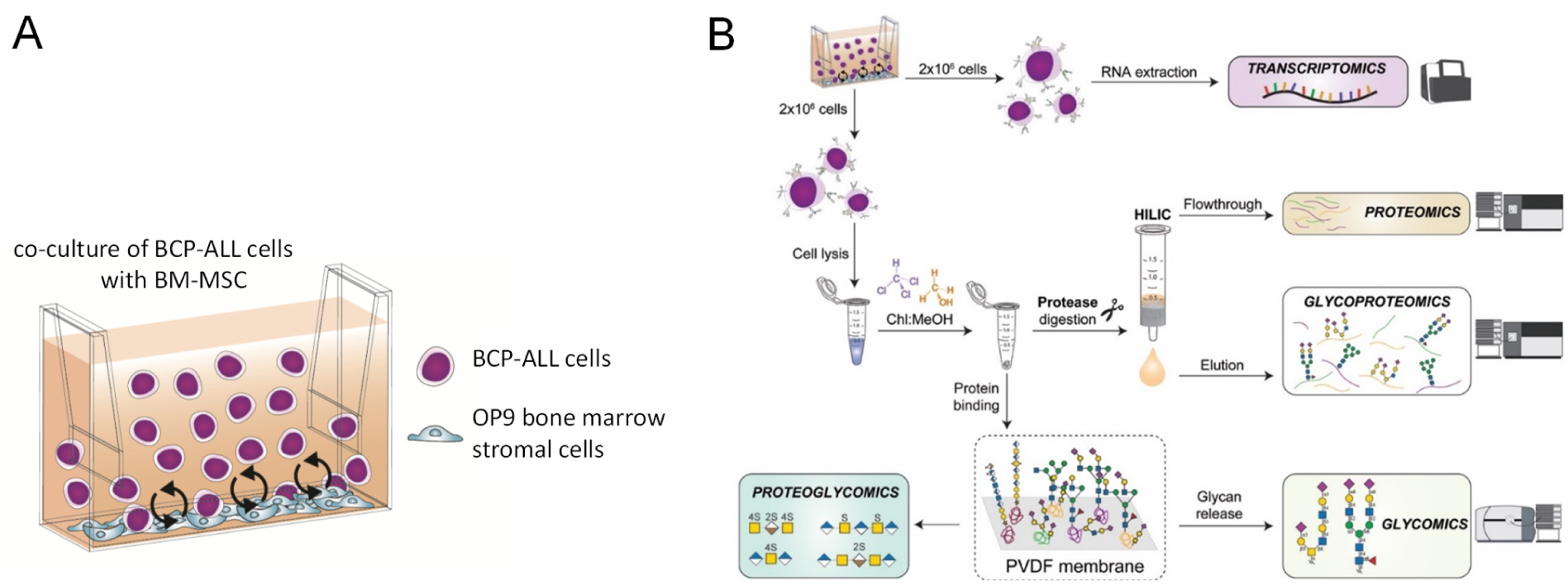
Schematic overview of Omics analysis of drug insensitivity development of human BCP-ALL cells treated with the chemotherapy drug vincristine. (**A**) Schematic illustration of co-culture of leukemia cells with stromal support. Leukemia cells traffic dynamically between the stromal support layer and the medium; cells were harvested from the supernatant for omics analysis (**B**) Schematic outline of omics analysis.

**Figure S2.**
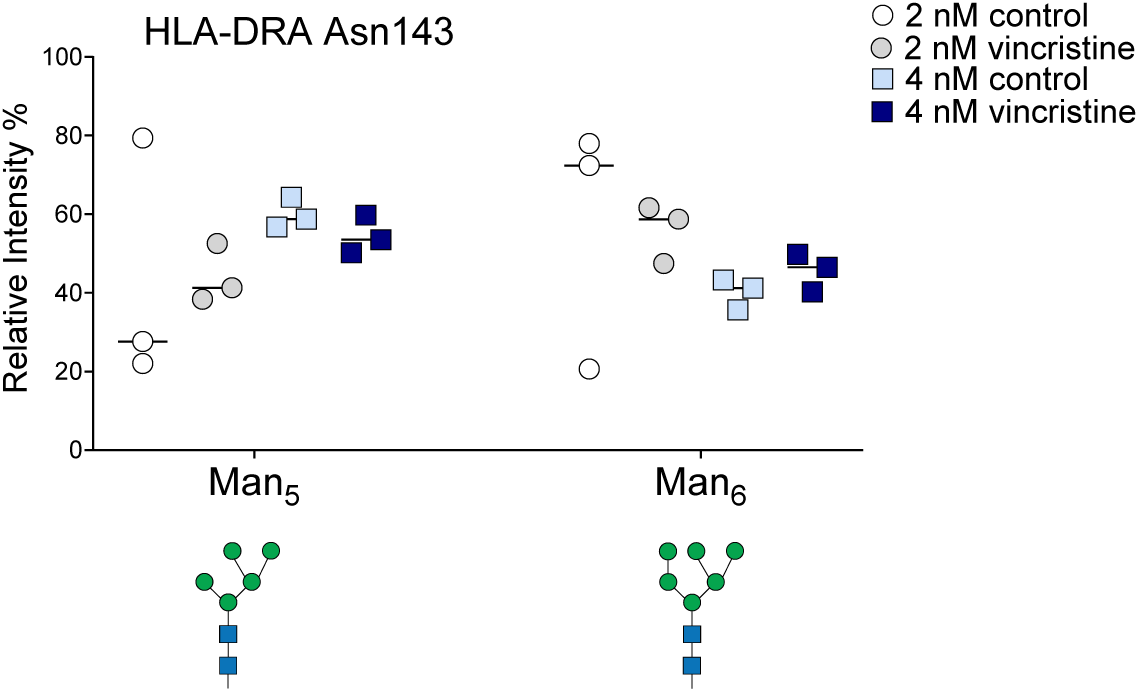
Relative intensity of N-glycans attached to HLA-DRA Asn143.

**Figure S3.**
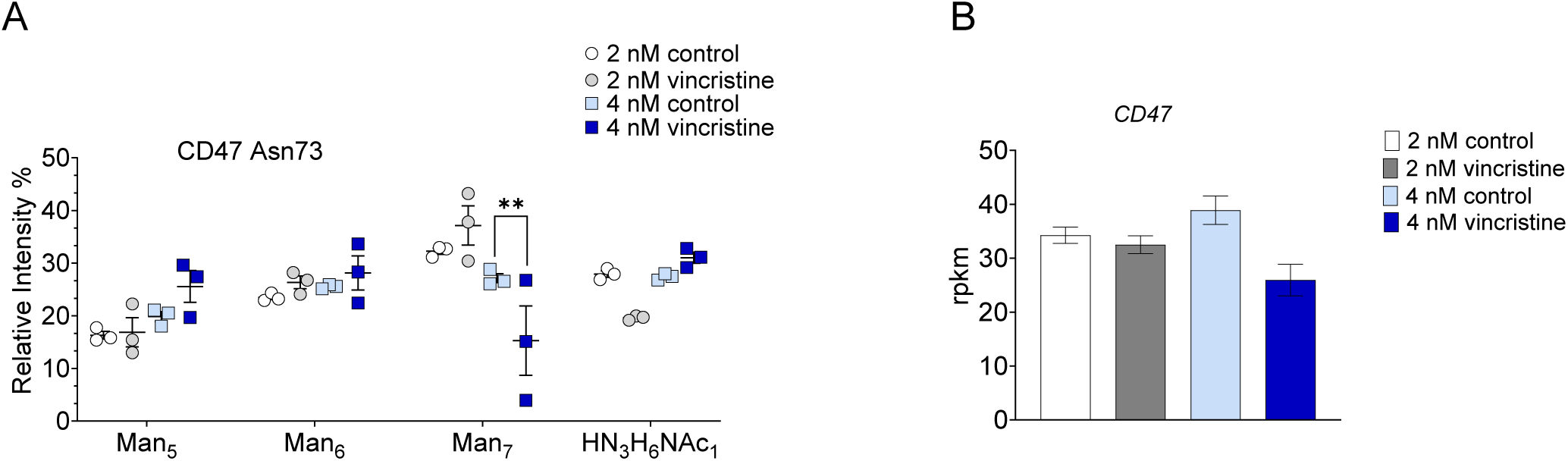
CD47 glycosylation and expression. (**A**) Asn73 N-linked glycosylation. (**B**) RNA expression. **<p<0.01

**Figure S4.**
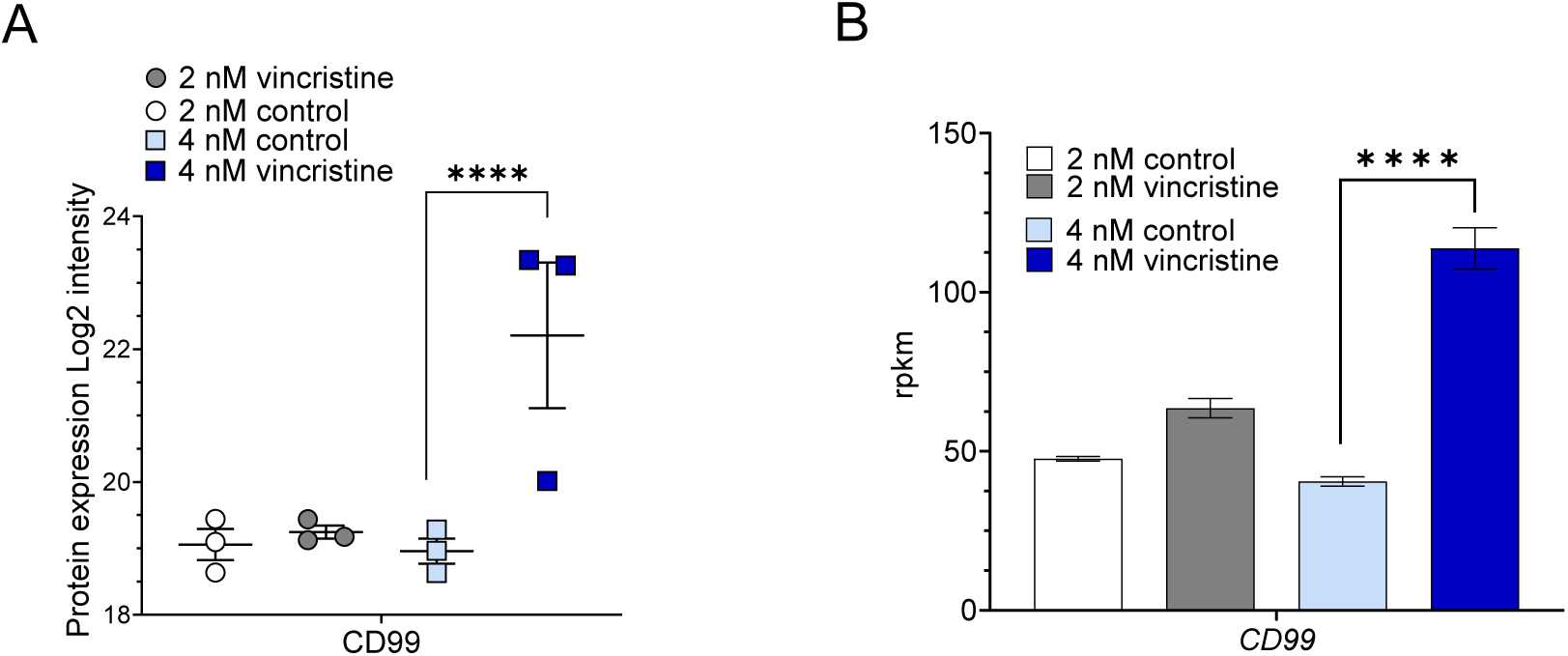
Increased protein and mRNA expression of CD99 in d30 DTP-BCP-ALL cells. (**A**) log2 intensity of protein expression (**B**) RNA levels in ICN13 cells treated as indicated. ****p<0.001

**Figure S5.**
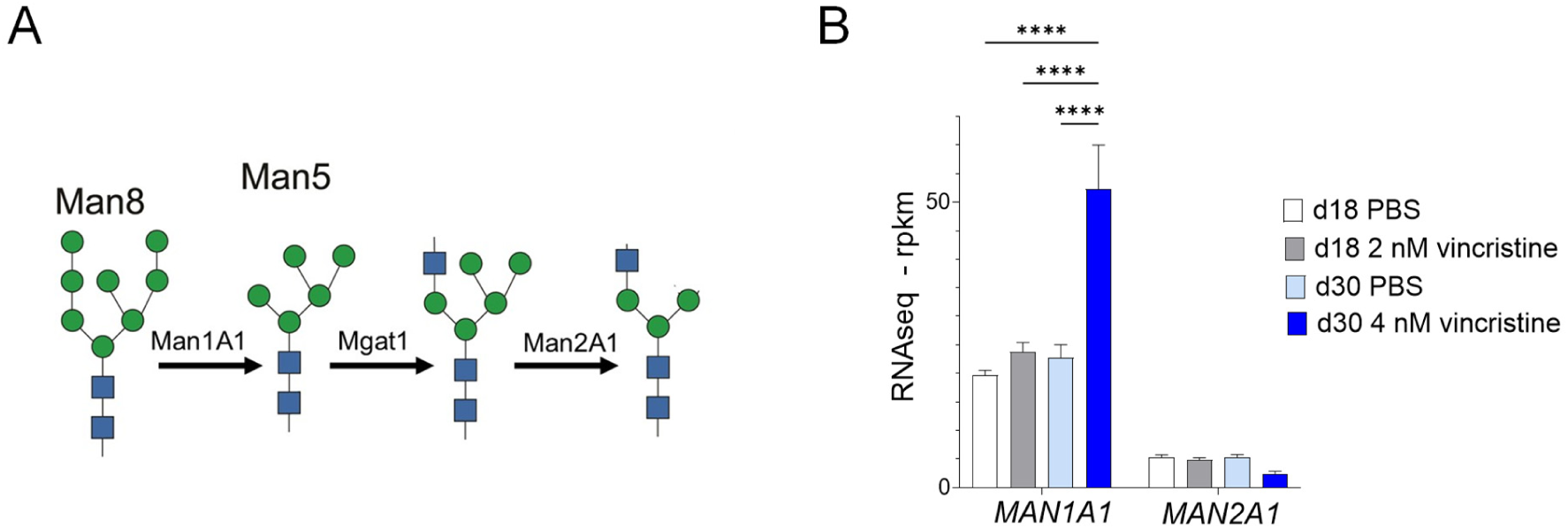
Oligomannose synthesis. (**A**) structures/enzymes (**B**) mRNA levels in the indicated conditions. ****p<0.001

